# Skap2 Regulates Atherosclerosis through Macrophage Polarization and Efferocytosis

**DOI:** 10.1101/857649

**Authors:** Danielle Hyatt, Allison E. Schroeder, Ashita Bhatnagar, David E. Golan, Kenneth D. Swanson, Francis J. Alenghat

## Abstract

**Rationale:** Atherosclerosis causes more deaths than any other pathophysiologic process. It has a well-established inflammatory, macrophage-mediated component, but important and potentially protective intracellular macrophage processes in atherosclerosis remain enigmatic. Src Kinase-Associated Phosphoprotein 2 (Skap2) is a macrophage-predominant adaptor protein critical for cytoskeletal reorganization, and thereby, for macrophage migration and chemotaxis. The role of macrophage Skap2 in atherosclerosis is unknown and deserves exploration.

**Objective:** To establish the critical role of Skap2 in macrophage-mediated atherosclerotic plaque homeostasis.

**Results:** In human arterial gene expression analysis, *Skap2* expression is enriched in macrophage-containing areas of human atheroma, and the transcript level varies with plaque characteristics. We have discovered that deletion of *Skap2* accelerates atherosclerosis by threefold in *ApoE*^-/-^ mice on standard diet. Skap2 expression is switched on only as monocytes differentiate into macrophages, so *Skap2*^-/-^ monocytes have no defect in infiltrating the atheroma. On the other hand, once they fully differentiate, Skap2-deficient macrophages cannot polarize efficiently into alternatively-activated, regulatory cells, and instead they preferentially polarize toward the classical pro-inflammatory phenotype both *ex vivo* and within the developing atheroma. This defect extends to polarized effector functions, as *ex vivo* analysis of macrophage phagocytosis of dying foam cells indicates that Skap2 is required for the regulatory process of efferocytosis.

**Conclusions:** Taken together, our findings support a model in which Skap2 drives a regulatory, efferocytic mode of behavior to quell atherosclerosis.

**CONDENSED ABSTRACT / SUMMARY:** Skap2—a macrophage protein found in the human atheroma—is atheroprotective. Skap2-null mice, whose foam cells do not migrate well due to a defect in integrin-induced cytoskeletal rearrangement, have accelerated atherosclerosis. Skap2 is not expressed in monocytes but becomes important once they reach the atheroma and become macrophage foam cells, at which point it drives toward a regulatory, anti-inflammatory polarization state required for efficient efferocytosis of dying foam cells. Thus, Skap2 drives a protective, regulatory mode of behavior, supporting the fact that macrophages are not solely deleterious in atherosclerosis, and further pointing to efferocytosis as a target for therapy.

There are no relationships to disclose.

## INTRODUCTION

Vascular disease due to atherosclerosis accounts for more mortality that any other pathophysiologic process. Common sites of clinically significant atherosclerosis include the coronary arteries, carotid arteries, aorta, and other major branches arising from the aorta. Atherosclerotic lesions begin as accumulations of lipoproteins within the intima of the artery, often at sites of arterial stress or disrupted flow patterns. In response to oxidized lipoprotein-elicited cytokine expression, circulating monocytes adhere to the endothelium and are recruited to the intima where they differentiate into macrophages and take up inflammatory lipoprotein particles to become foam cells. Foam cells within the growing lesions induce more oxidative stress, protease activity, and pro-inflammatory cytokine secretion. In response to these stimuli, smooth muscle cells migrate and divide within the intima, and cycles of fibrosis, proliferation, and apoptosis ensue within the growing plaque (1). As atherosclerosis progresses in this manner, lesion growth into the arterial lumen may be sufficient to impair blood flow and induce distal ischemia. Additionally, plaques may become unstable and prone to rupture, which would result in thrombus formation and distal infarction. While much attention has been paid to the role of macrophages in atherogenesis, it is not fully understood how critical aspects of macrophage effector and homeostatic functions, including the balance between inflammatory and regulatory macrophages populations within the atherosclerotic lesion, affect processes involved in this pathophysiology.

Src Kinase-Associated Phosphoprotein 2 (Skap2), an adaptor protein encoded by the gene *Skap2*, is normally only expressed in lymphoid and myeloid cells. Among these cells, its presence appears most critical for macrophage function (2-6). In macrophages, Skap2 binds to the transmembrane Signal regulatory protein α (Sirpα) and the adapter protein ADAP (5), and it plays a central role transmitting signals from ligand-bound integrins to direct actin cytoskeletal rearrangement required for cell spreading and migration (7). Given the importance of macrophage movement in regulating inflammation, understanding how Skap2 impacts atheroma formation would point to novel insights that could inform the development of targeted treatments for inflammatory pathways governing atherosclerosis.

The Apolipoprotein E (*ApoE*) knockout mouse is an established and practical model of atherosclerosis. On a standard diet, ApoE-deficient mice develop atherosclerotic lesions that progress with age (8,9). We therefore sought to test the effect of *Skap2* deletion on atherosclerosis in *ApoE*^-/-^ mice. Although Skap2 is necessary for the transmission of macrophage integrin outside-in signaling, it was unclear whether Skap2-deficiency would exacerbate or ameliorate the atherosclerotic phenotype. On one hand, impairment of monocyte-to-macrophage differentiation or blocking macrophage proliferation ameliorates atherogenesis. For instance, atherosclerosis is significantly attenuated in mice that lack cytokines and chemokines that guide monocyte or macrophage proliferation and recruitment, such as macrophage colony stimulating factor (M-CSF), monocyte chemotactic protein-1 (MCP-1), or MCP-1’s receptor (10-12). Likewise, inhibiting macrophage interaction with CD40L or depleting macrophages of cell adhesion molecules, including integrins, also reduces lesion severity (13-15). Knockout of the pro-apoptotic and anti-proliferative p53 yields more severe lesions in *ApoE*^-/-^ mice, suggesting that uncontrolled macrophage proliferation is atherogenic (16,17). Still, while such past findings demonstrate that macrophages can contribute to formation and progression of lesions, it is also possible that macrophages play protective roles by clearing the arterial intima of potentially damaging lipoproteins or cellular debris, and that clinically significant atherosclerosis could represent a dysregulation or overloading of this system. Here, we find that Skap2 regulates the inflammatory response in atherosclerosis by promoting regulatory (or so-called “M2”) macrophage polarization and effector functions.

## METHODS

### Antibodies and Reagents

Antibodies against-Skap2 (12926), CD206 (18704-1-AP), and β-actin (66009-1-Ig) were obtained from Proteintech Group. Antibodies against F4/80 (123109), CD3 (100201), CD34 (119301), and CD86 (105001) were obtained from Biolegend. AcLDL (L23380), anti-F4/80 (MF48000), and Goat anti-Rat IgG (H+L) Secondary Antibody, Biotin-XX conjugate (A10517) were obtained from Invitrogen. LPS was from Sigma, INFγ and IL4 were from ProSpec.

### Human Atheroma Skap2 Expression

Human Skap2 and CD68 expression was obtained from the Gene Expression Omnibus (GEO) data repository (18). Expression was analyzed in human atherosclerotic aorta samples with matched comparisons to non-atherosclerotic internal mammary artery in coronary artery bypass patients (NCBI GEO database accession GSE40231) (19), and in human carotid endarterectomy samples comparing plaque to adjacent areas free of macroscopic disease (NCBI GEO database accession GSE43292) (20). The Skap2 analysis was also done in laser-dissected macrophages from ruptured vs. stable vs. fibrocalcific carotid plaques NCBI GEO database accession GSE41571) (21).

### Mice, Cell Culture and Macrophage Polarization

*Skap2*^*-/-*^ C57BL/6 mice and *ApoE*^*-/-*^ C57BL/6 mice (Jackson Laboratories) were described previously (8,22,23). Mice bearing these respective alleles were intercrossed, and backcrossed onto *ApoE*^*-/-*^ C57BL/6 for a minimum of six generations. Mice were maintained at the Harvard and University of Chicago animal facilities under pathogen-free conditions and were sacrificed at 10-24 weeks. All procedures were approved by the Harvard and University of Chicago Institutional Animal Care and Use Committees.

Bone marrow-derived macrophages (BMMs) were differentiated *ex vivo* for 7 days as described previously (24). These adherent macrophages were cultured in Dulbecco’s Modified Eagles Medium (DMEM) containing 10% FBS, 1% Penicillin/Streptomycin, and 10% macrophage colony stimulating factor (M-CSF) containing medium (CMG-condition media) (25). Bone-marrow derived monocytes (BMMos) were defined as the non-adherent viable cells differentiated *ex vivo* for 7 days in this medium (26). Polarization of BMMs was initiated after the 7 days of differentiation (M0) using this medium with 5 ng/ml LPS and 12 ng/ml interferon-γ for 24 hours vs. 10 ng/ml IL-4 for 48 hours to generate inflammatory (“M1”) vs. regulatory (“M2’) polarized macrophages, respectively, as previously described (27). All cells were maintained at 37**°**C in 5% CO_2_.

### Reverse Transcriptase Quantitative PCR

RNA was isolated from non-polarized and polarized macrophages using the GeneJET RNA purification kit (Thermo Scientific), and quantified using Nanodrop (Thermo Scientific). cDNA was reverse transcribed using iScript Supermix (Biorad) and subjected to qPCR using the following primers:*Ym1*: F: 5’ CAGGTCTGGCAATTCTTCTGAA 3’ R: 5’ GCTTGCTCATGTGTGTAAGTGA 3’, *Arg1*: F: 5’ CTCCAAGCCAAAGTCCTTAGAG 3’ R: 5’ AGGAGCTGTCATTAGGGACATC 3’, *Nos2*: F: 5’ GTTCTCAGCCCAACAATACAAGA 3’ R: 5’ GTGGACGGGTCGATGTCAC 3’, *Tnfa*: F: 5’ CCCTCACACTCAGATCATCTTCT 3’ R: 5’ GCTACGACGTGGGCTACAG 3’ and *ß-actin*: F: 5’ GGCTGTATTCCCCTCCATCG 3’ R: 5’ CCAGTTGGTAACAATGCCATGT 3’. Detection was achieved using advanced universal SYBR green supermix (Biorad) by qPCR in a CFX96 thermal cycler (Biorad).

### Western blot analysis

Cultured cells were lysed using 1% Triton X-100, 50mM Tris-HCl, and 150mM NaCl and clarified by centrifugation. Protein was quantified using bicinchoninic acid protein assay (Pierce). The lysate proteins were resolved using SDS-PAGE, transferred to nitrocellulose, and blocked in TBST with 2% milk for 1 hour. Blots were probed with the indicated primary antibodies for 1 hour and detected using horseradish peroxidase-conjugated secondary antibodies. Bound antibodies were detected using SuperSignal West Femto ECL substrate (34095, Thermo Scientific) and imaged using ChemiDoc and iLab software (Biorad).

### Cell Selection and FACS

F4/80^+^ BMMs and BMMos were selected using magnetic beads (M-1002-020; Solulink). The cells were blocked in FACS buffer (200mM EDTA and 1% FBS in PBS) for 30 minutes and then incubated with rat anti-mouse F4/80 primary antibody for 1 hour. The cells were then washed, blocked in FACS buffer and incubated with biotin anti-rat antibody for 1 hour. F4/80^+^ cells were then collected using streptavidin conjugated magnetic beads prior to FACS preparation. For analysis of adherent cells, they were detached using 5mM EDTA in PBS for 5 minutes at 37**°**C prior to FACS preparation. The cells were washed and resuspended in FACS buffer for detection of surface markers or fixed using 2% paraformaldehyde and then washed, permeabilized, and resuspended in FACS buffer for detection of intracellular proteins. For staining, cells were blocked with FACS buffer for 1 hour on ice, incubated with primary antibodies in FACS buffer for 1 hour, washed with FACS buffer, blocked for 45 minutes, and incubated with secondary antibodies conjugated to the indicated fluorophore for 45 minutes in FACS buffer. Data were acquired using an LSRII 4-12 Flow cytometer (BD Biosciences) and analyzed using Flowing analytical software.

### Microscopy

Tissue sections were frozen in OCT at -80**°**C. Sections were cut at 10 μm for immunofluorescence analysis and at 16 μm for Oil Red O analysis. Tissue and cells were fixed in 2% PFA, 25mM PIPES pH 6.8, 129mM KCl, 20% sucrose, 5mM EDTA. For immunofluorescence of intracellular proteins, sections were permeabilized using 0.1% Triton X-100 in PBS, pH 7.4. Cells and tissue sections were blocked using 2% BSA for 45 minutes, incubated with primary antibodies using 2% BSA at room temperature for 1 hour, washed in PBS, blocked using 2% BSA for 30 minutes, and incubated in secondary antibodies using 2% BSA at room temperature for 45 minutes. Tissue sections and cells were imaged using a Zeiss Axioskop microscope.

### AcLDL Uptake and Efferocytosis Assays

BMMs were incubated at 37**°**C with Alexa-488-conjugated AcLDL (0.5-1.0 μg/ml) for 30 minutes to produce foam cells. The cells were then washed with PBS and serum free-DMEM was added. The cells were exposed to 5 minutes of UV-irradiation and then placed at 37**°**C for 36 hours to induce apoptosis. The apoptotic cells were detached and placed back at 37**°**C for 24 hours. These apoptotic foam cells were then incubated with viable WT and Skap2^-/-^ BMMs for 30-120 minutes in a 1:1 ratio. The cells were washed with PBS and stained with PE-conjugated F4/80. Images were analyzed using ImageJ software (NIH).

### Foam Cell Migration Assay

BMMs were plated at confluence on petri dishes for 16 hours, and 1 μg/ml AcLDL was then added for 30 minutes. The foam cells were then washed once with PBS. Scratch wounds were introduced into the cultures using a fine straight edge cell scraper, and then washed sufficiently to remove non-adherent cells. Fresh macrophage growth medium was added and cultures were incubated at 37**°**C. Images were taken at 0 and 8 hours and quantified for cell density within the scratch using ImageJ.

### Statistics

Statistical testing of differences was performed using the 2-tailed Student’s t-test.

## RESULTS

### Skap2 is expressed in human atheroma and varies with plaque characteristics

*Skap2* expression was analyzed in different sites of atherosclerosis obtained from humans using the Gene Expression Omnibus (GEO) data repository (18). Compared to non-diseased internal mammary artery (19), Skap2 expression was generally higher in atherosclerotic areas or aorta sampled during coronary artery bypass grafting surgery (**Figure 1A**). It is important to note that, in order to permit proper saphenous vein grafting, the portion of the aorta sampled in this study cannot be severely diseased, suggesting that Skap2 appears in early stages of lesion formation. In human carotid endarterectomy samples, plaques were compared to adjacent areas free of macroscopically visible disease (20), and again, *Skap2* expression is generally higher in diseased artery (**Figure 1B**). In these samples, consistent with Skap2 as a macrophage-specific protein, *Skap2* expression correlated directly with the expression of human macrophage marker CD68 (**Figure 1C**), and importantly, this correlation exists not only in the visible plaque but also in the adjacent artery free of macroscopic disease, again supporting early lesion progression with macrophages in some of those areas. The Skap2 analysis was also done in laser dissected macrophages from ruptured vs. stable vs. fibrocalcific carotid artery (21), demonstrating that the degree of Skap2 expression varies with plaque characteristics and stability, with the most Skap2 found in stabilized, non-calcified plaques (**Figure 1D**)

**Figure 1.**
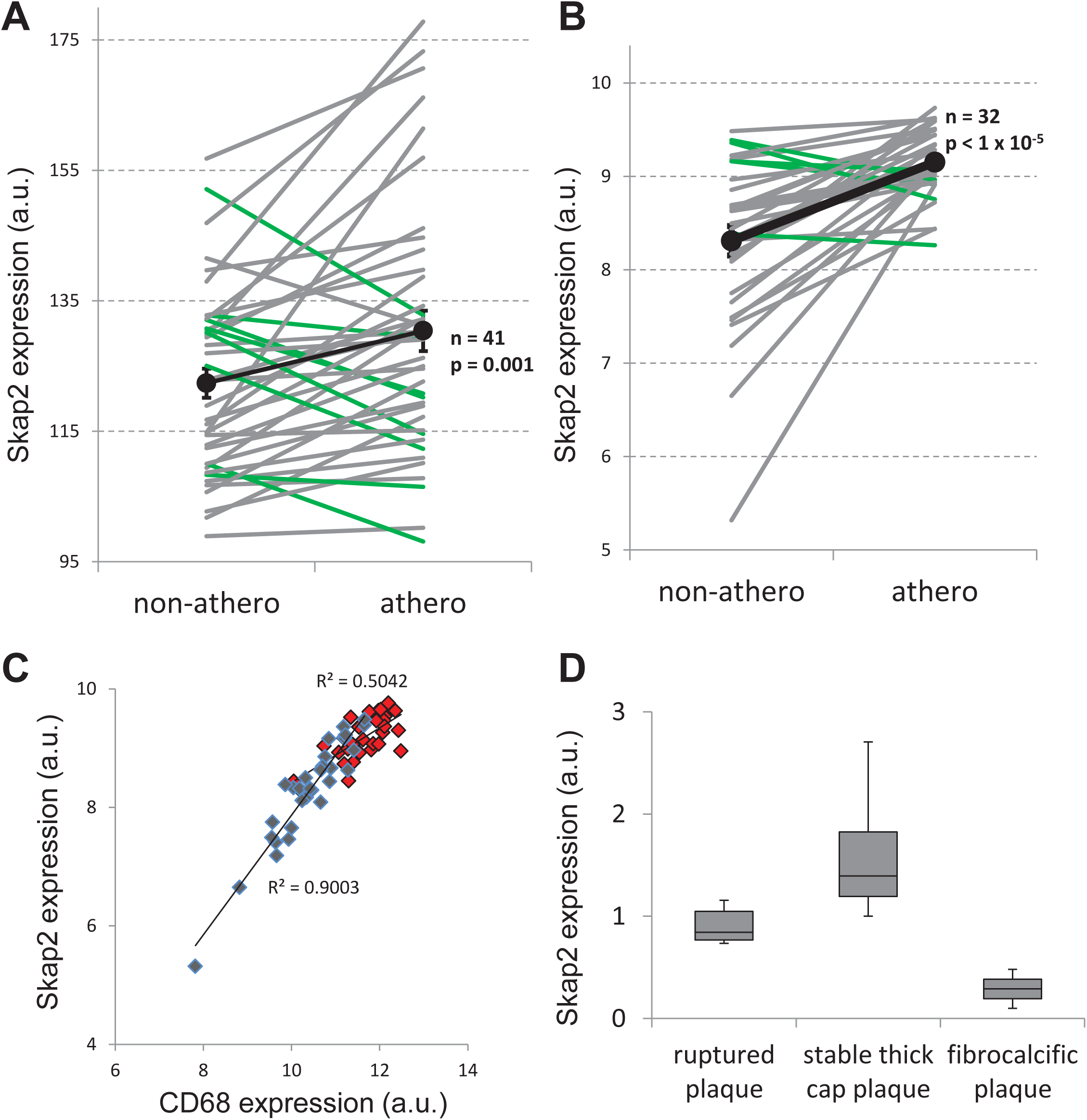
Skap2 is expressed in human atheroma and varies with plaque characteristics. (**A**) Skap2 transcript expression is increased in most human atheroma samples from aorta (gray) compared to matched internal mammary artery free of macroscopic disease. A minority of patients had less Skap2 transcript in the atheroma (green). (**B**) Skap2 expression is increased in most human carotid atheroma samples compared to matched adjacent carotid free of macroscopic disease. In both **A** and **B**, means are shown in black. (**C**) Arterial Skap2 expression correlates with macrophage content (as reflected by CD68 expression) in artery without visible disease (blue) and in artery with macroscopic disease (red). The correlation in arteries without visible disease suggests early lesion progression with macrophages. While Skap2 and CD68 expression are higher in arteries with visible disease, their correlation is less strong, suggesting it may vary with plaque characteristics. (**D**) Skap2 expression varies by plaque characteristics, upregulated in stable plaques and reduced in calcific and ruptured plaque. Data shown as box-and-whisker plot.

### *Skap2* deletion exacerbates atherosclerosis

To probe whether Skap2 is involved in atherosclerosis, we generated and maintained Skap2-replete and Skap2-deficient mice on an *ApoE*-knockout background and fed them a standard diet. At 24 weeks, *Skap2*^*-/-*^*/ApoE*^*-/-*^ mice exhibited approximately 3-fold greater plaque burden within their aortic roots compared to *Skap2*^*+/-*^*/ApoE*^*-/-*^ (**Figure 2A**). This 3-fold difference was also apparent at 18-weeks (**Figure 2B**). There were no significant differences in weight (**Figure 2C**) or in total cholesterol, HDL, and LDL (**Figure 2D**) between the *Skap2*-knockout and *Skap2*-expressing mice in these cohorts.

**Figure 2.**
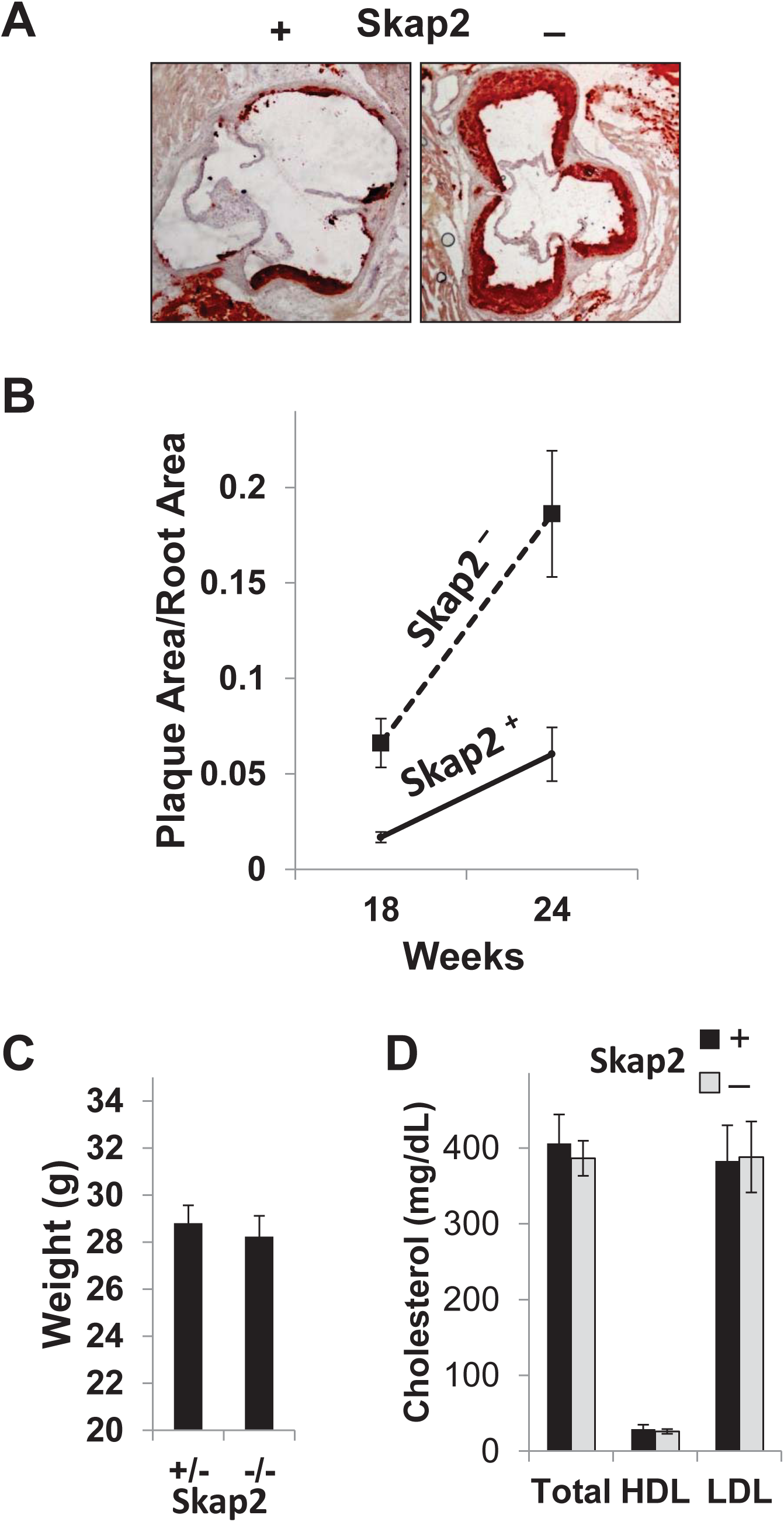
Skap2-deficient ApoE^-/-^ mice have accelerated atherosclerosis. (**A**) 24-week-old Skap2^-/-^/ApoE^-/-^ mice exhibited accelerated atherosclerotic plaque formation in the aortic roots compared to Skap2^+/-^/ApoE^-/-^ littermates visualized by Oil Red O staining. (**B**) The increased plaque size is approximately 3-fold greater in Skap2^-/-^/ApoE^-/-^ 24-week-old mice. Significant differences were also seen at 18 weeks. (**C**-**D**) There were no differences in weight or lipid profiles between the Skap2^-/-^/ApoE^-/-^ and Skap2^+/-^/ApoE^-/-^ mice.

Skap2 did not impact T cell or endothelial cell distribution within the atheromata, as measured by CD3 and CD34 immunohistochemical staining, respectively (**Figure 3A**). Despite the requirement for Skap2 for efficient macrophage cytoskeletal rearrangement, chemotaxis, and migration *in vitro* (7,28), the atheromata of Skap2-deficient mice recruit abundant macrophages as indicated by F4/80 immunohistochemical and immunofluorescence staining, and the macrophage density is higher in Skap2-deficient lesions compared to Skap2-replete lesions (**Figure 3A-C**). Thus, Skap2 does not appear to impair monocytic recruitment into the developing lesions.

**Figure 3.**
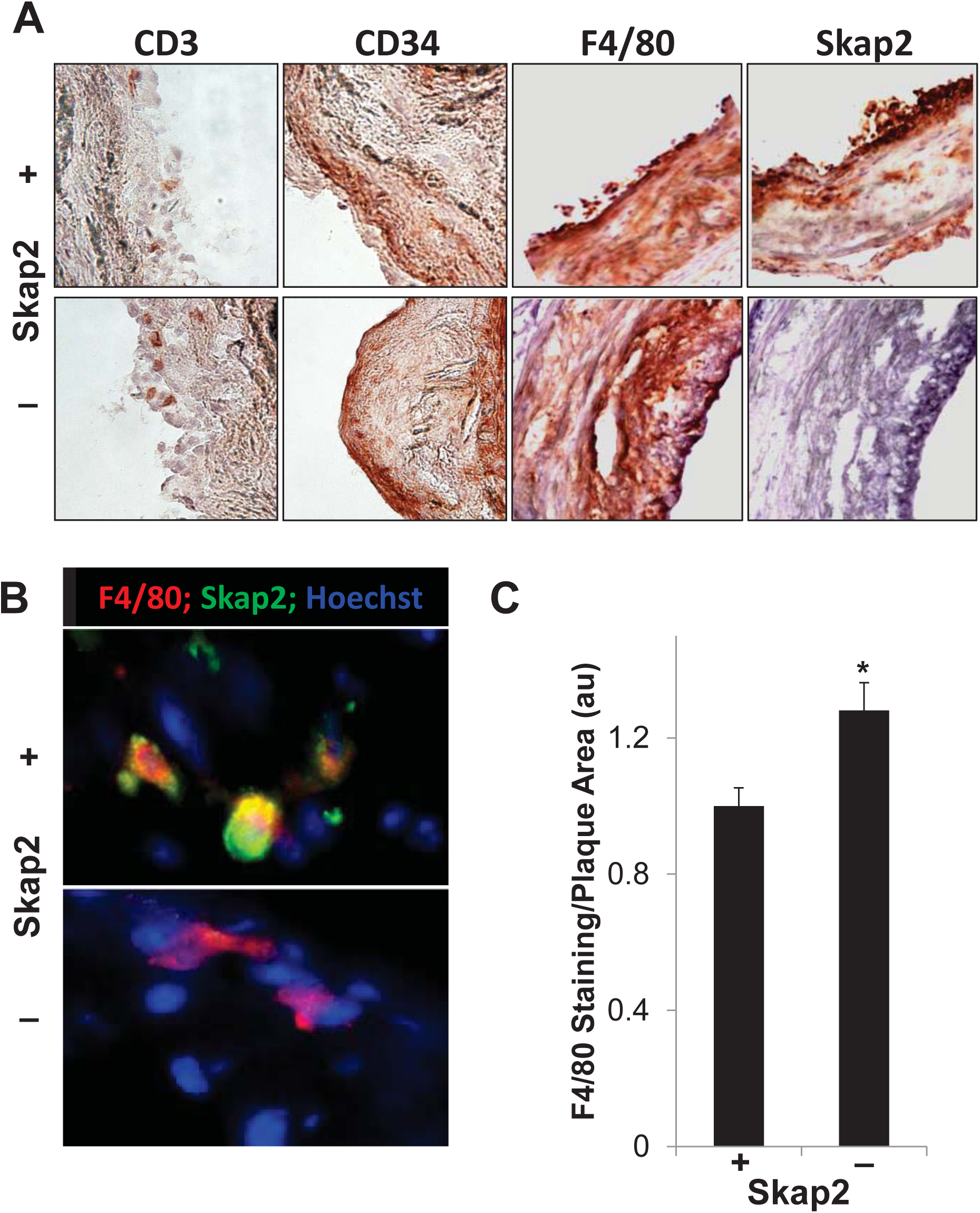
Within the atheroma, Skap2 deficiency has no visible effect on cells other than macrophages. (**A**) CD3 and CD34 staining of atherosclerotic plaques from Skap2^-/-^/ApoE^-/-^ and Skap2^+^ApoE^-/-^ mice revealed similar distributions of both T cell and vascular endothelial cells, respectively. Macrophage content (F4/80) was present as assessed by IHC (24-week mice, **A**) and immunofluorescence staining (18-week mice, **B**), so Skap2-deficient macrophages show no defect incorporating into the atheroma, and with greater macrophage density in Skap2^-/-^ mice (**C**). * p < 0.05, n=9 and 13 for Skap2^+^ and Skap2^-^, respectively.

### Skap2 in the myeloid population

Given the increased atheroma size and lesional macrophage content in the absence of Skap2, we investigated the overall inflammatory status of *Skap2*^*-/-*^*/ApoE*^*-/-*^ mice by measuring a panel of common cytokines in the serum of 24 week old mice. IL1α, IL6, and IL8 were significantly elevated in *Skap2*-deficient mice, suggestive of a macrophage-centric pro-inflammatory state (**Figure 4A**). The levels of other circulating cytokines tested (IL2, IL4, IL5, IL10, IL12, IL13, TNFα, IFNγ) were not altered by *Skap2* deletion (not shown).

**Figure 4.**
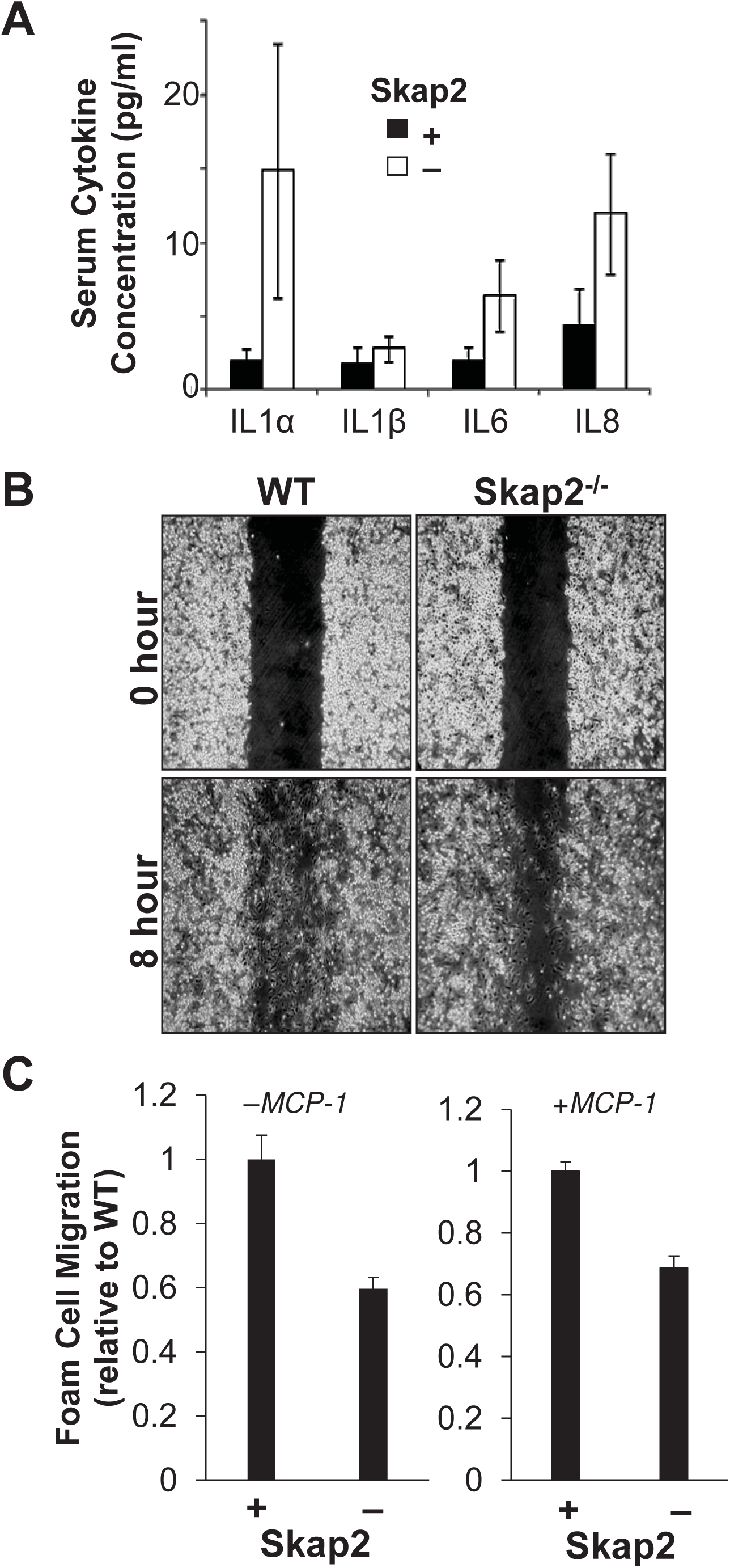
Skap2-deficiency elicits a macrophage centric pro-inflammatory signature and diminished foam cell migration. (**A**) Circulating levels of IL1α, IL6, and IL8 are elevated in Skap2-deficient, ApoE^-/-^ mice. No differences in IL2, IL4, IL5, IL10, IL12, IL13, IFNγ and TNFα levels were observed between Skap2^-/-^/ApoE^-/-^ and Skap2^+/+^/ApoE^-/-^ mice. Foam cells were produced from BMM by inducing uptake of 1 μg/ml acetylated LDL. (**B**) Scratches were introduced into confluent populations of foam cells (upper panels); after 8 hours, Skap2-deficient foam cells displayed impaired migration (lower panels), quantified both with and without 50 ng/ml MCP-1 in the media (**C**). *p < 0.5, n = 6.

We have previously shown that *Skap2*^-/-^ macrophages do not migrate at normal rates (7). Given that foam cells are abundant in the *Skap2*^*-/-*^ atheroma, we sought to dissect the potential contribution of Skap2 in foam cell macrophage migration and found that *Skap2*^*-/-*^ AcLDL-laden foam cells demonstrated impaired migration in a standard scratch assay (**Figure 4B and 4C**). This impairment persisted in the presence of MCP-1 (**Figure 4C**). The requirement of Skap2 for foam cell migration suggests that, although macrophage incorporation into lesions is not impacted, intralesional macrophage behavior is governed by this protein.

Skap2 is expressed and functional in the myeloid lineage (lymphoid cells rely on a closely related protein Skap1) (3,29-31). However, the relative importance of Skap2 in monocytes versus terminally differentiated macrophages is not known and could shed light on why *Skap2*-null cells readily home to sites of vascular inflammation (32,33). Consistent with its role in cytoskeletal rearrangement, we find that Skap2 expression coincides with the adhesion and spreading events that characterize monocyte maturation to macrophage. Indeed, we found that upon culturing bone marrow with M-CSF to drive macrophage differentiation *ex vivo*, Skap2 levels increased from an undetectable level in early cultured cells, to heterogeneous, very low levels in mature monocytes, to high levels in adherent macrophages (**Figure 5A and 5B**). Furthermore, F4/80-positive bone marrow-derived monocytes (BMMos), the immediate non-adherent precursors to BMMs, exhibit far less Skap2 than the macrophages they become; specifically, BMMs expressed six-fold more Skap2 compared to BMMos, and a large fraction of F4/80^+^ BMMos do not express appreciable Skap2 (**Figure 5C**). These findings further support that fully differentiated macrophages/foam cells depend on Skap2 but their monocyte precursors do not.

**Figure 5.**
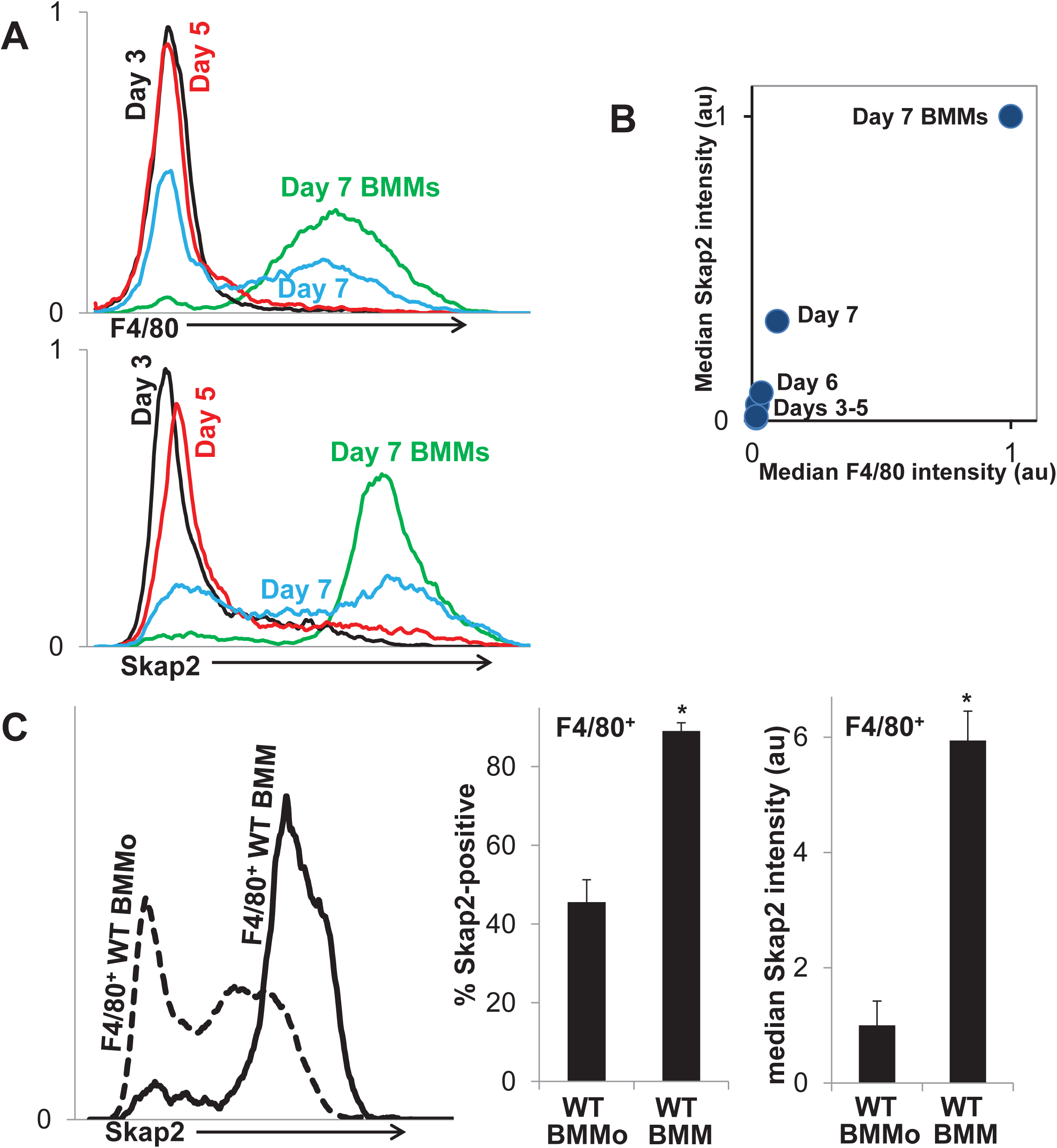
Skap2 is highly expressed in macrophages but not in their monocyte precursors. Skap2 expression increases during macrophage differentiation from bone marrow cells in culture with incubation in M-CSF-containing media (**A**-**B**). Macrophage differentiation was determined by F4/80 staining and adherence. FACS analysis of these cultures revealed that F4/80^high^ macrophages express higher levels of Skap2 than pre-macrophage populations in the culture, with the most pronounced increase in expression occurring with the transition from monocyte (BMMo, day 7) to macrophage (BMM). Early monocytes express negligible Skap2. Even amongst the F4/80^+^ cells, BMMs have a ∼6-fold increase in median Skap2 intensity compared to monocytes (**C**). Skap2 expression increases significantly upon adherence, as monocytes become macrophages. * p < 0.05.

### Skap2 is required for expression of M2 polarization markers

Based on the macrophage-predominant expression of Skap2, the requirement of Skap2 for foam cell migration, and the macrophage-centric inflammatory cytokine signature in Skap2-deficient mice, we further probed whether Skap2 impacts the inflammatory phenotype of the macrophage. We polarized WT BMMs, denoted M0 for the non-polarized phenotype, toward the inflammatory (so-called “M1”) or the regulatory (“M2”) phenotype using LPS and IFNγ or IL4, respectively (27). Regulatory macrophages have greater Skap2 expression than other macrophages, and, in the absence of Skap2, expression of the regulatory macrophage surface marker CD206 is reduced (**Figure 6A**). Immunofluorescence staining of cells also demonstrated that regulatory macrophages expressed Skap2 at higher levels than either M0 or inflammatory macrophages (**Figure 6B**).

**Figure 6.**
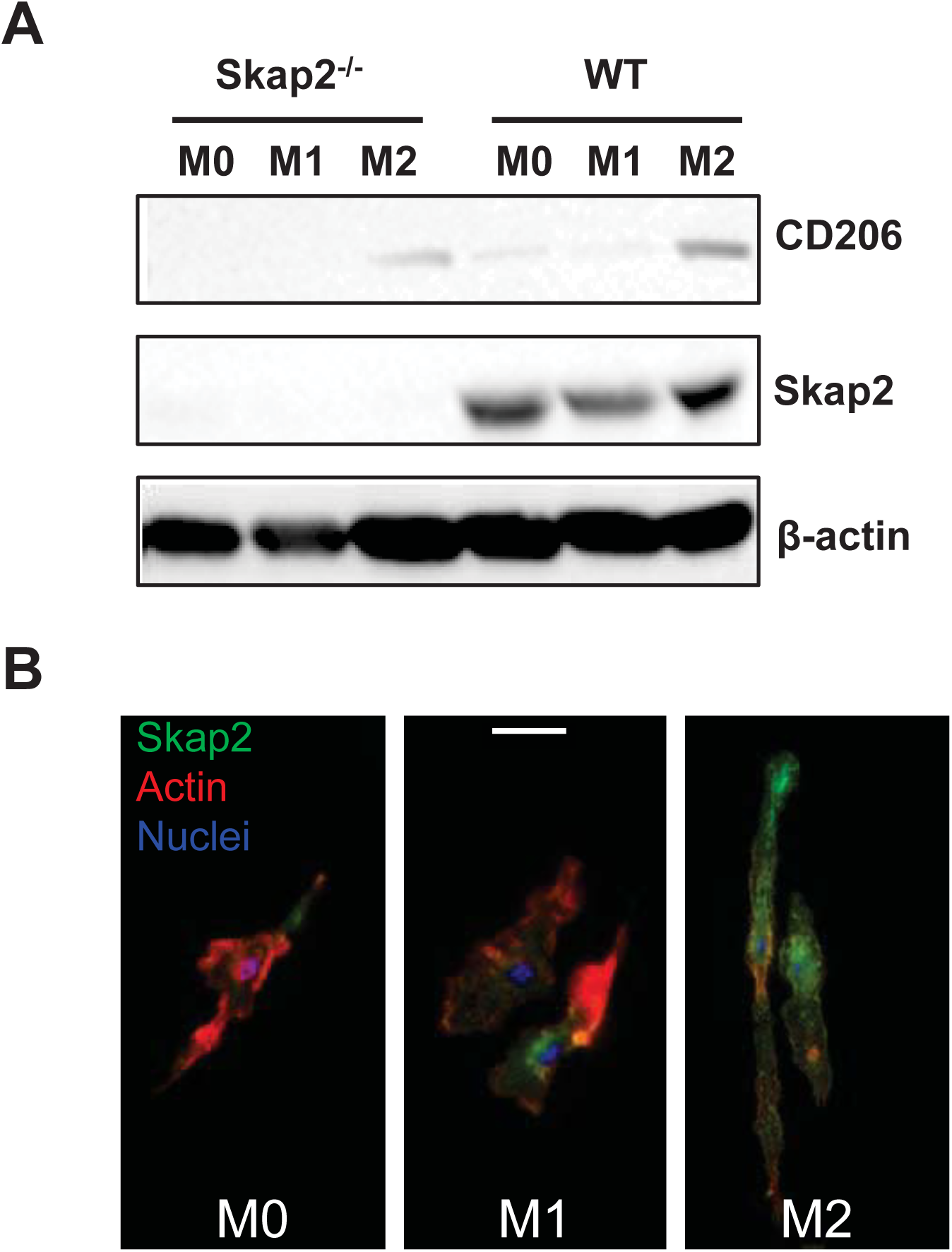
The expression of Skap2 is accentuated in regulatory macrophages and is required for M2 surface marker expression. (**A**) Regulatory “M2” macrophages express more Skap2 than BMMs (M0) and pro-inflammatory “M1” macrophages. Compared to WT M2 macrophages, Skap2^-/-^ M2 macrophages express lower levels of the M2 marker CD206. (**B**) Increased Skap2 expression (green) in regulatory macrophages is recapitulated using immunofluorescence staining in WT polarized macrophages co-stained for actin with phalloidin (red). Scale bar = 30μm.

Given the increased expression of Skap2 in regulatory macrophages, we asked whether Skap2 played a role in inflammatory versus regulatory macrophage polarization. BMMs were polarized towards either type of macrophage and analyzed for canonical M1 markers, *Tnfα* and *Nos2*, and M2 markers, *Arg1* and *Ym1* to validate the polarization culture system (**Figure 7A**) (27,34). Skap2^-/-^ inflammatory macrophages (M1), compared to WT macrophages, expressed higher mRNA levels of the established M1 markers. At the same time, Skap2^-/-^ regulatory macrophages (M2) expressed lower levels of the M2 markers compared to WT (**Figure 7B**).

**Figure 7.**
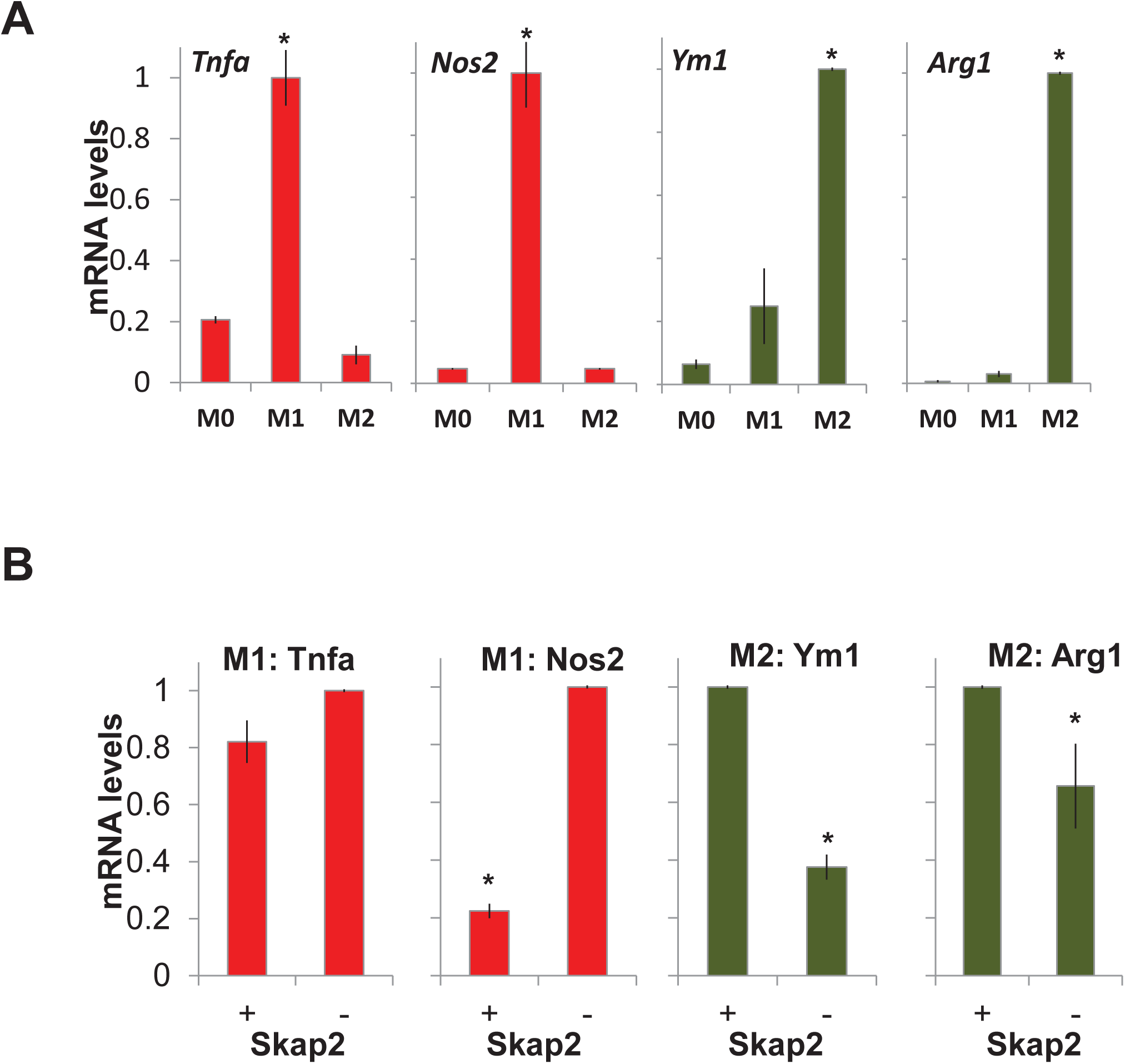
Skap2 promotes regulatory macrophages as measured by transcription of polarization markers. BMMs were polarized *in vitro* to M1 and M2 macrophages. (**A**) qPCR was used to analyze the expression of canonical transcript markers for M1 (Tnfα and Nos2) and M2 macrophages (Ym1 and Arg1). (**B**) In this system, compared to polarized wild-type (+) cells, Skap2^-/-^ M1 macrophages express augmented levels of inflammatory M1 markers and Skap2^-/-^ M2 macrophages express depressed regulatory M2 markers. Relative mRNA levels are normalized to the highest value per plot, *p < 0.05.

To explore whether this disruption of the balance between inflammatory and regulatory polarized states extends to the atheroma, we stained for M1/M2 surface markers in aortic root lesions of 18 week and 24 week old *Skap2*^*-/-*^*/ApoE*^*-/-*^ mice. Staining with the M1 marker CD86 and the M2 marker CD206 revealed a two-fold increase in the M1:M2 ratio within the atherosclerotic lesions of the Skap2^-/-^ mice (**Figure 8A and 8B**). Thus, while *Skap2*-null monocytes can reach the developing lesion, the resulting intralesional macrophages skew toward a more inflammatory signature.

**Figure 8.**
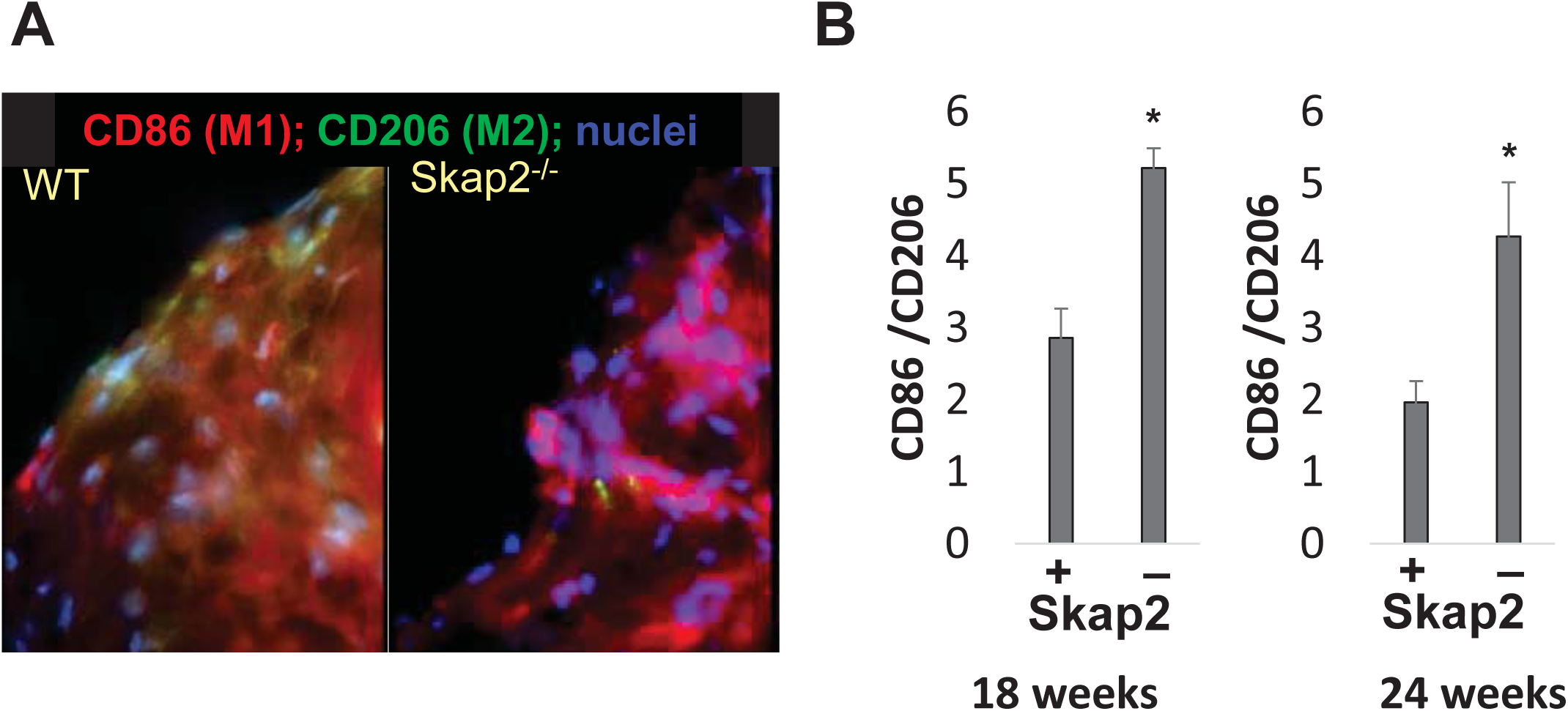
Skap2 is associated with regulatory polarization in atherosclerosis. (**A**) Analysis of lesions within the aortic roots of Skap2^-/-^/ApoE^-/-^ mice revealed an increased ratio of M1 macrophages (marked by CD86) versus M2 macrophages (CD206) relative to that seen in lesions from Skap2^+/+^/ApoE^-/-^ mice. (**B**) This skewing of the ratio of CD86-to-CD206 was evident in Skap2 deficient atheromata at both 18 and 24 weeks. n = 9, *p < 0.05.

### Skap2 is required for efficient efferocytosis

Skap2 deficiency reduces the levels of actin-rich ruffles on the surface of macrophages (7) suggesting that there may be a defect in fluid endocytic function in Skap2 mutant BMMs. However, using three classical markers—Lucifer yellow, FITC-Dextran, and Alexa 488-AcLDL— we found no significant differences in non-receptor-mediated micropinocytosis, multilectin receptor-mediated uptake, or scavenger receptor-mediated uptake, respectively (**Figure 9A**) (35). Given that the clearance of apoptotic debris arising from the high turnover of foam cells within the lesion is an important function performed by alternatively-activated, regulatory macrophage populations during atherogenesis, we measured efferocytosis in WT and *Skap2*^*-/-*^ BMM cultures. To measure macrophage efferocytosis *in vitro*, we loaded Alexa-488-conjugated AcLDL into BMMs and subsequently exposed them to brief UV-irradiation followed by serum starvation to induce apoptosis. The resulting apoptotic foam cells were incapable of adhering or spreading on glass substrates like normal BMMs, but retained their nuclei and general structure. When these dying foam cells were introduced to viable BMMs, *Skap2*^*-/-*^ BMMs displayed inefficient efferocytosis of this cellular debris (**Figure 9B and 9C**). This central component of clearance of inflammatory debris appears to require Skap2, supporting its role in maintaining macrophage-mediated intralesional homeostasis.

**Figure 9.**
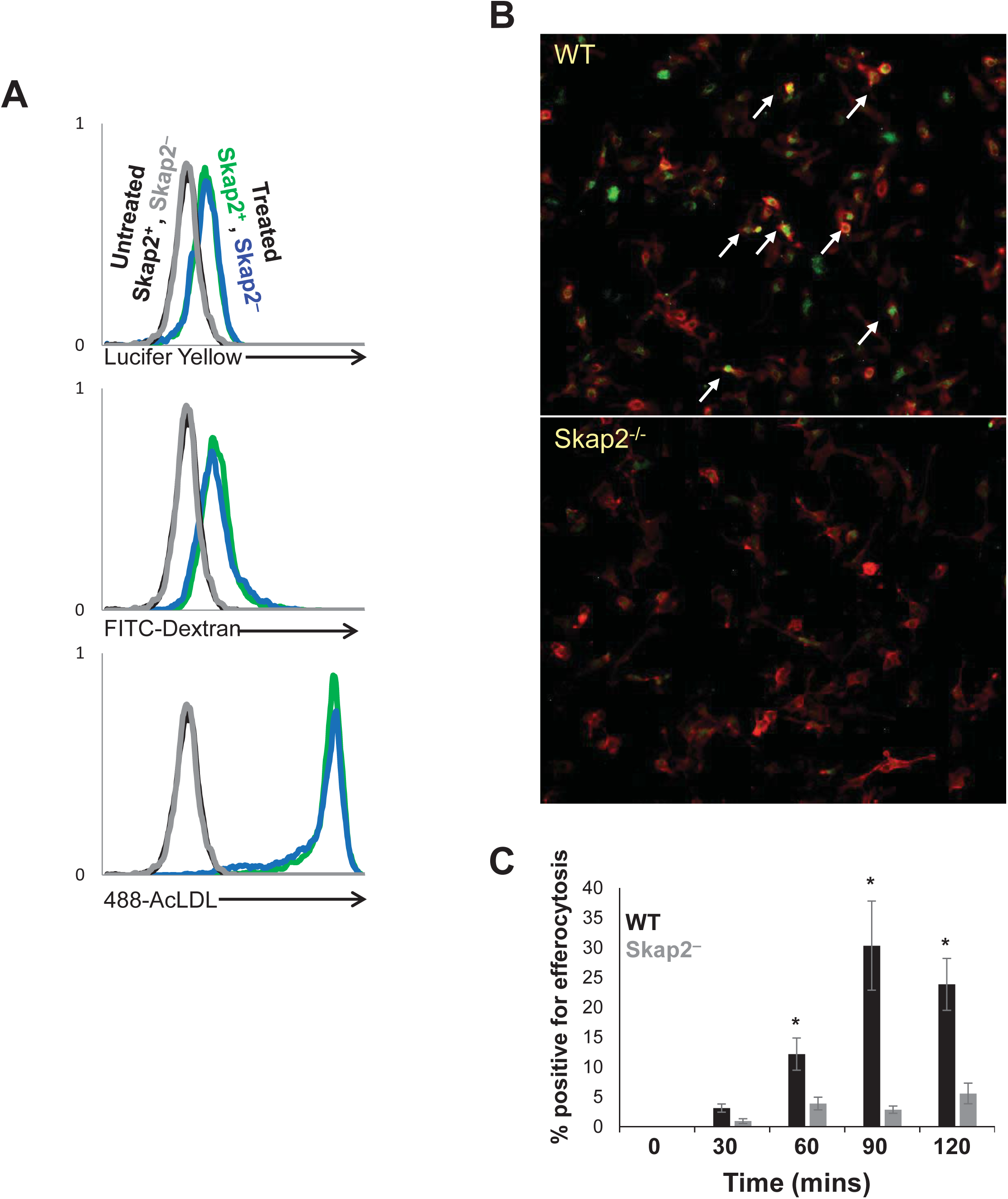
Skap2 deficiency does not alter standard endocytic pathways but does impact efferocytosis. (**A**) FACS analysis shows that nonspecific macropinocytosis of Lucifer Yellow, mannose receptor-mediated uptake of FITC-Dextran and scavenger receptor-mediated uptake of AcLDL is not impacted by Skap2 deficiency. (**B**-**C**) In contrast, when BMMs were co-incubated with apoptotic FITC-AcLDL loaded foam cells, Skap2^-/-^ BMMs inefficiently efferocytosed these cells, a process which peaked at 90 minutes of co-incubation in WT cells.

## DISCUSSION

Our findings demonstrate that Skap2 is important in anti-inflammatory processes that act during atherogenesis. *Skap2* expression increases with macrophage differentiation and its very low level of expression in monocytes suggests that they do not require Skap2 to function in circulation. Thus, there is preserved macrophage content of *Skap2*^-/-^ lesions, which are three-fold larger than Skap2-replete lesions in atherosclerotic mice. Once macrophages without Skap2 incorporate into the lesion, however, there are important defects in the maintenance of intralesional homeostasis to keep inflammation in check. Our findings support a model in which Skap2 is required for efficient foam cell migration, proper polarization to the regulatory macrophage mode, and participation in the process of efferocytic debris clearance.

Macrophage polarization within the atherosclerotic plaque has emerged as a critical aspect of lesion homeostasis (36,37). Classically activated inflammatory macrophages drive pro-inflammatory responses through the production of inflammatory cytokines and reactive oxygen species whereas alternatively activated regulatory macrophages secrete anti-inflammatory cytokines and lipid mediators (38). *Skap2* deletion leads to dysregulation of macrophage polarization resulting in the production of more M1-like (pro-inflammatory) macrophages demonstrated both *in vitro* and in the aortic roots of Skap2^-/-^/ApoE^-/-^ mice. Furthermore, defective regulatory macrophage differentiation in Skap2 deficient animals, together with impaired foam cell migration and inefficient efferocytosis of apoptotic foam cells, strongly suggests a model in which Skap2 plays a prominent role in promoting these anti-inflammatory effector functions to tamp down on inflammatory processes within developing atheromata. *In vitro* and mouse models of polarized macrophages show that classically-activated macrophages are involved in inflammatory plaque progression, whereas alternatively-activated macrophages resolve this inflammation (34,39). Our findings indicate that Skap2 helps to maintain balance between these opposing phenotypes. In many pathophysiologic processes, regulatory alternatively-activated macrophages help to resolve inflammation and orchestrate repair by synthesizing mediators, such as TGFβ and vascular endothelial growth factor, central to tissue remodeling (40). Preferential expression of Skap2 in regulatory macrophages suggests that the protein is more important in maintaining this polarization state, and the protein and transcript marker expression pattern of Skap2-deficient macrophages support this.

The clearance of apoptotic cells by efferocytosis is a crucial aspect of successful resolution of inflammatory processes (41). Although the mechanisms of macrophage foam cell death in athero-prone regions is not well understood, several studies have shown that macrophage apoptosis increases as lesions progress, and thus, clearance of the resulting cellular debris is increasingly necessary for either resolution of inflammation or limiting the inflammation-induced damage occurring during plaque progression (42,43). Growth factor deprivation, the presence of toxic cytokines, increased ER stress and excess accumulation of lipoprotein-derived cholesterol by macrophages all may contribute to the increased apoptosis at later stages of atherosclerosis (44,45). In the absence of efficient apoptotic cell uptake, dying cells can undergo secondary necrosis. This autolytic process intensifies the inflammatory response and leads to further formation of the necrotic core that can destabilize the plaque (42,46,47). Regulatory macrophages are more efficient at efferocytosis than the classically-activated inflammatory macrophages (48,49). *Skap2*^*-/-*^ BMMs exhibited reduced efferocytosis of apoptotic foam cells in our *ex vivo* system, demonstrating intrinsic impairment in efferocytosis. Combined with overall augmented inflammatory and diminished regulatory macrophage polarization, this may account for a significant part of the increase in plaque burden in Skap2-deficient mice.

The way in which Skap2 drives efferocytosis is unclear. Apoptotic cells can be recognized by the tethering receptor TIM4, and it has been demonstrated that TIM4-mediated phagocytosis is dependent on integrin-induced Src family kinase activation (50). We have previously established that Skap2 acts within a pathway the drives cytoskeletal rearrangement downstream of integrins binding to their extracellular matrix. Src family kinase activation in response to integrins is dependent on Skap2 expression (7). Therefore, the defect in efferocytosis in Skap2^-/-^ BMM could be attributed to compromised integrin signaling and Src family kinase activation downstream of TIM4 engagement with phosphatidyl serine on the surface of apoptotic cells. However, it is clear that efferocytosis is distinct from other phagocytic processes, so other explanations are quite possible. Indeed, no defect in either Fc-or complement mediated phagocytosis was detected in these macrophages (22), suggesting differences in Skap2 requirements for these pathways. Because Skap2 binds to Sirpα (5,7), one intriguing possibility is that Skap2 impacts the interaction between CD47 and Sirpα (the so-called “don’t eat me” signal) to promote efferocytosis by preventing anti-phagocytosis (51,52); recent work has demonstrated that blocking CD47 signaling to macrophages can enhance efferocytosis and reduce atherosclerosis (53), so future work should focus on how CD47-blockade and Skap2 signaling interact.

## CONCLUSIONS

Taken together, our data demonstrate that Skap2 keeps inflammatory macrophage/foam cell phenotypes and processes in check. It is required for the differentiation of regulatory M2 macrophage populations and plays a counter-regulatory function in the production of the macrophage inflammatory phenotype. By controlling such homeostasis within the atheroma, Skap2 therefore appears to regulate atherosclerosis by driving macrophages toward a regulatory, efferocytic phenotype. We hope these findings motivate future work to understand the mechanisms by which Skap2 and its signaling partners (such as Sirpα) determine the shifting balance between inflammatory and anti-inflammatory modes to point to rational new approaches for treatment of atherosclerosis and other chronic inflammatory diseases.

## PERSPECTIVES

### COMPETENCY IN MEDICAL KNOWLEDGE

Macrophages can serve a protective role in early atherosclerosis and participate in a process of inflammation resolution and clearance through efferocytosis. Promoting pro-resolving macrophage phenotypes and stimulating plaque debris clearance hold promise as potential future preventive therapies for atherosclerosis.

### TRANSLATIONAL OUTLOOK

Future translational studies should examine targeting macrophage polarization and efferocytosis in human macrophages from patients with atherosclerosis, and subsequently employ therapies such as CD47/Sirpα blockade, which acts through Skap2, to enhance efferocytosis in an effort to drive plaque stabilization or regression.

## ABBREVIATIONS

ApoE: Apolipoprotein E
BMM: Bone marrow-derived macrophage
BMMo: Bone marrow-derived monocyte
GEO: Gene Expression Omnibus
M-CSF: Macrophage colony stimulating factor
MCP-1: Monocyte chemotactic protein-1
Sirpα: Signal regulatory protein α
Skap2: Src Kinase-Associated Phosphoprotein 2

## ACKNOWLEDGEMENTS

Microscopy was performed at the Imaging Center at University of Chicago and the Nikon Imaging Center at Harvard Medical School. D.H. performed the research, analyzed the data, and wrote the paper. D.E.G., and K.D.S. helped design research and assisted in writing the paper. F.J.A. designed research, performed research, analyzed data, and wrote the paper.

---

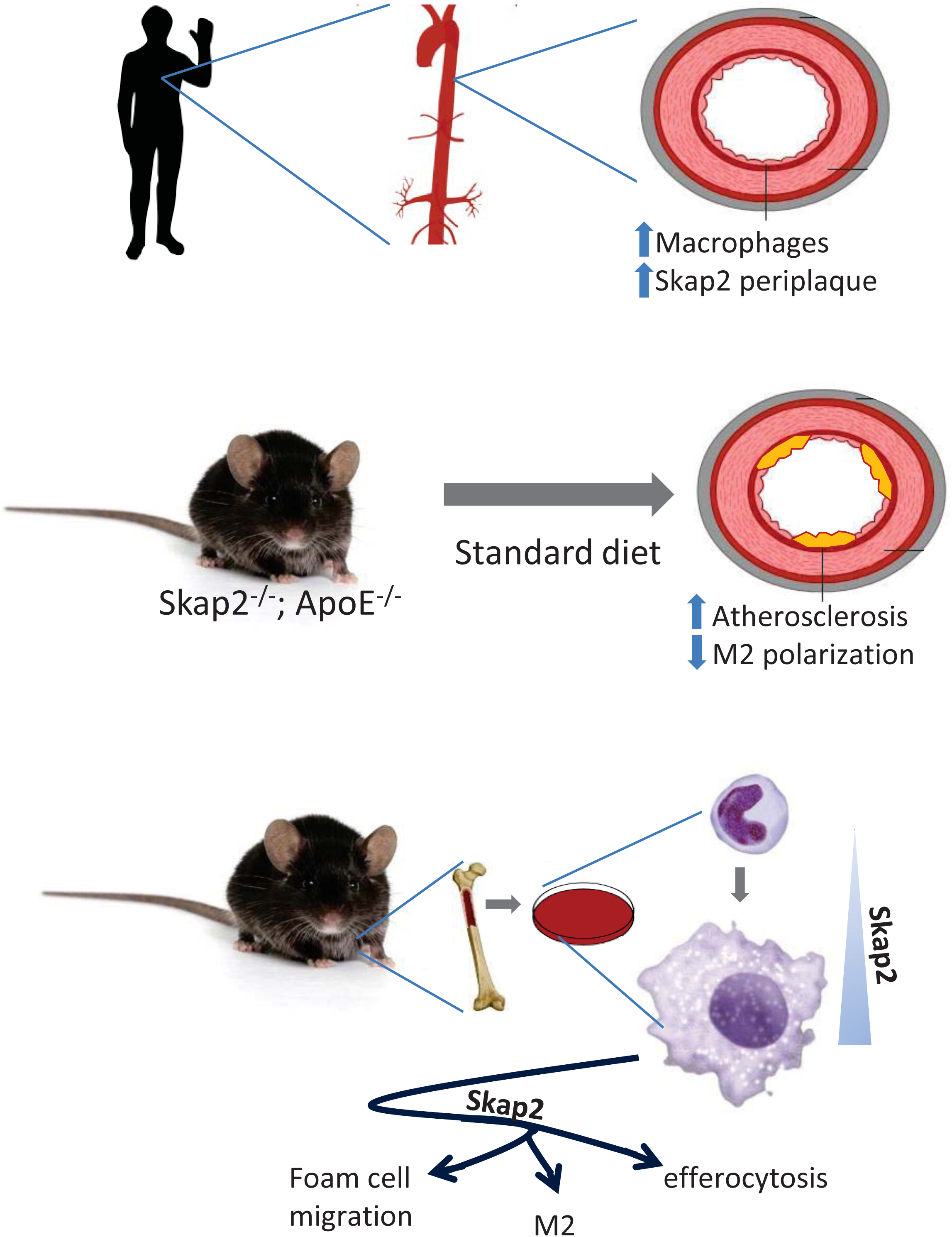

## REFERENCES

1. Libby P. Inflammation in atherosclerosis. Nature 2002;420:868–74.

2. Curtis DJ, Jane SM, Hilton DJ, Dougherty L, Bodine DM, Begley CG. Adaptor protein SKAP55R is associated with myeloid differentiation and growth arrest. Exp Hematol 2000;28:1250–9.

3. Kouroku Y, Soyama A, Fujita E, Urase K, Tsukahara T, Momoi T. RA70 is a src kinase-associated protein expressed ubiquitously. Biochem Biophys Res Commun 1998;252:738–42.

4. Bourette RP, Therier J, Mouchiroud G. Macrophage colony-stimulating factor receptor induces tyrosine phosphorylation of SKAP55R adaptor and its association with actin. Cell Signal 2005;17:941–9.

5. Timms JF, Swanson KD, Marie-Cardine A et al. SHPS-1 is a scaffold for assembling distinct adhesion-regulated multi-protein complexes in macrophages. Curr Biol 1999;9:927–30.

6. Black DS, Marie-Cardine A, Schraven B, Bliska JB. The Yersinia tyrosine phosphatase YopH targets a novel adhesion-regulated signalling complex in macrophages. Cell Microbiol 2000;2:401–14.

7. Alenghat FJ, Baca QJ, Rubin NT et al. Macrophages require Skap2 and Sirpalpha for integrin-stimulated cytoskeletal rearrangement. J Cell Sci 2012.

8. Zhang SH, Reddick RL, Piedrahita JA, Maeda N. Spontaneous hypercholesterolemia and arterial lesions in mice lacking apolipoprotein E. Science 1992;258:468–71.

9. Whitman SC. A practical approach to using mice in atherosclerosis research. Clin Biochem Rev 2004;25:81–93.

10. Smith JD, Trogan E, Ginsberg M, Grigaux C, Tian J, Miyata M. Decreased atherosclerosis in mice deficient in both macrophage colony-stimulating factor (op) and apolipoprotein E. Proc Natl Acad Sci U S A 1995;92:8264–8.

11. Boring L, Gosling J, Cleary M, Charo IF. Decreased lesion formation in CCR2-/-mice reveals a role for chemokines in the initiation of atherosclerosis. Nature 1998;394:894–7.

12. Gu L, Okada Y, Clinton SK et al. Absence of monocyte chemoattractant protein-1 reduces atherosclerosis in low density lipoprotein receptor-deficient mice. Mol Cell 1998;2:275–81.

13. Mach F, Schonbeck U, Sukhova GK, Atkinson E, Libby P. Reduction of atherosclerosis in mice by inhibition of CD40 signalling. Nature 1998;394:200–3.

14. Zirlik A, Maier C, Gerdes N et al. CD40 ligand mediates inflammation independently of CD40 by interaction with Mac-1. Circulation 2007;115:1571–80.

15. Nageh MF, Sandberg ET, Marotti KR et al. Deficiency of inflammatory cell adhesion molecules protects against atherosclerosis in mice. Arterioscler Thromb Vasc Biol 1997;17:1517–20.

16. van Vlijmen BJ, Gerritsen G, Franken AL et al. Macrophage p53 deficiency leads to enhanced atherosclerosis in APOE*3-Leiden transgenic mice. Circ Res 2001;88:780–6.

17. Guevara NV, Kim HS, Antonova EI, Chan L. The absence of p53 accelerates atherosclerosis by increasing cell proliferation in vivo. Nat Med 1999;5:335–9.

18. Edgar R, Domrachev M, Lash AE. Gene Expression Omnibus: NCBI gene expression and hybridization array data repository. Nucleic Acids Res 2002;30:207–10.

19. Hagg S, Skogsberg J, Lundstrom J et al. Multi-organ expression profiling uncovers a gene module in coronary artery disease involving transendothelial migration of leukocytes and LIM domain binding 2: the Stockholm Atherosclerosis Gene Expression (STAGE) study. PLoS Genet 2009;5:e1000754.

20. Ayari H, Bricca G. Identification of two genes potentially associated in iron-heme homeostasis in human carotid plaque using microarray analysis. J Biosci 2013;38:311–5.

21. Lee K, Santibanez-Koref M, Polvikoski T, Birchall D, Mendelow AD, Keavney B. Increased expression of fatty acid binding protein 4 and leptin in resident macrophages characterises atherosclerotic plaque rupture. Atherosclerosis 2013;226:74–81.

22. Togni M, Swanson KD, Reimann S et al. Regulation of in vitro and in vivo immune functions by the cytosolic adaptor protein SKAP-HOM. Mol Cell Biol 2005;25:8052–63.

23. Piedrahita JA, Zhang SH, Hagaman JR, Oliver PM, Maeda N. Generation of mice carrying a mutant apolipoprotein E gene inactivated by gene targeting in embryonic stem cells. Proc Natl Acad Sci U S A 1992;89:4471–5.

24. Tushinski RJ, Oliver IT, Guilbert LJ, Tynan PW, Warner JR, Stanley ER. Survival of mononuclear phagocytes depends on a lineage-specific growth factor that the differentiated cells selectively destroy. Cell 1982;28:71–81.

25. Takeshita S, Kaji K, Kudo A. Identification and characterization of the new osteoclast progenitor with macrophage phenotypes being able to differentiate into mature osteoclasts. J Bone Miner Res 2000;15:1477–88.

26. Francke A, Herold J, Weinert S, Strasser RH, Braun-Dullaeus RC. Generation of mature murine monocytes from heterogeneous bone marrow and description of their properties. J Histochem Cytochem 2011;59:813–25.

27. Becker L, Liu NC, Averill MM et al. Unique proteomic signatures distinguish macrophages and dendritic cells. PLoS One 2012;7:e33297.

28. Swanson KD, Tang Y, Ceccarelli DF et al. The Skap-hom dimerization and PH domains comprise a 3’-phosphoinositide-gated molecular switch. Mol Cell 2008;32:564–75.

29. Liu J, Kang H, Raab M, da Silva AJ, Kraeft SK, Rudd CE. FYB (FYN binding protein) serves as a binding partner for lymphoid protein and FYN kinase substrate SKAP55 and a SKAP55-related protein in T cells. Proc Natl Acad Sci U S A 1998;95:8779–84.

30. Marie-Cardine A, Verhagen AM, Eckerskorn C, Schraven B. SKAP-HOM, a novel adaptor protein homologous to the FYN-associated protein SKAP55. FEBS Lett 1998;435:55–60.

31. Jo EK, Wang H, Rudd CE. An essential role for SKAP-55 in LFA-1 clustering on T cells that cannot be substituted by SKAP-55R. J Exp Med 2005;201:1733–9.

32. Geissmann F, Manz MG, Jung S, Sieweke MH, Merad M, Ley K. Development of monocytes, macrophages, and dendritic cells. Science 2010;327:656–61.

33. Taghavie-Moghadam PL, Butcher MJ, Galkina EV. The dynamic lives of macrophage and dendritic cell subsets in atherosclerosis. Ann N Y Acad Sci 2014;1319:19–37.

34. Peled M, Fisher EA. Dynamic Aspects of Macrophage Polarization during Atherosclerosis Progression and Regression. Frontiers in immunology 2014;5:579.

35. Michael DR, Ashlin TG, Davies CS et al. Differential regulation of macropinocytosis in macrophages by cytokines: implications for foam cell formation and atherosclerosis. Cytokine 2013;64:357–61.

36. Moore KJ, Tabas I. Macrophages in the pathogenesis of atherosclerosis. Cell 2011;145:341–55.

37. Thorp E, Subramanian M, Tabas I. The role of macrophages and dendritic cells in the clearance of apoptotic cells in advanced atherosclerosis. Eur J Immunol 2011;41:2515–8.

38. Adamson S, Leitinger N. Phenotypic modulation of macrophages in response to plaque lipids. Curr Opin Lipidol 2011;22:335–42.

39. Brenner C, Franz WM, Kuhlenthal S et al. DPP-4 inhibition ameliorates atherosclerosis by priming monocytes into M2 macrophages. Int J Cardiol 2015;199:163–169.

40. Laskin DL, Sunil VR, Gardner CR, Laskin JD. Macrophages and tissue injury: agents of defense or destruction? Annu Rev Pharmacol Toxicol 2011;51:267–88.

41. Serhan CN, Brain SD, Buckley CD et al. Resolution of inflammation: state of the art, definitions and terms. FASEB J 2007;21:325–32.

42. Tabas I. Consequences and therapeutic implications of macrophage apoptosis in atherosclerosis: the importance of lesion stage and phagocytic efficiency. Arterioscler Thromb Vasc Biol 2005;25:2255–64.

43. Bhatia VK, Yun S, Leung V et al. Complement C1q reduces early atherosclerosis in low-density lipoprotein receptor-deficient mice. Am J Pathol 2007;170:416–26.

44. Tabas I. Apoptosis and plaque destabilization in atherosclerosis: the role of macrophage apoptosis induced by cholesterol. Cell Death Differ 2004;11 Suppl 1:S12–6.

45. Devries-Seimon T, Li Y, Yao PM et al. Cholesterol-induced macrophage apoptosis requires ER stress pathways and engagement of the type A scavenger receptor. J Cell Biol 2005;171:61–73.

46. Schrijvers DM, De Meyer GR, Herman AG, Martinet W. Phagocytosis in atherosclerosis: Molecular mechanisms and implications for plaque progression and stability. Cardiovasc Res 2007;73:470–80.

47. Silva MT. Secondary necrosis: the natural outcome of the complete apoptotic program. FEBS Lett 2010;584:4491–9.

48. Michlewska S, Dransfield I, Megson IL, Rossi AG. Macrophage phagocytosis of apoptotic neutrophils is critically regulated by the opposing actions of pro-inflammatory and anti-inflammatory agents: key role for TNF-alpha. FASEB J 2009;23:844–54.

49. McPhillips K, Janssen WJ, Ghosh M et al. TNF-alpha inhibits macrophage clearance of apoptotic cells via cytosolic phospholipase A2 and oxidant-dependent mechanisms. J Immunol 2007;178:8117–26.

50. Flannagan RS, Canton J, Furuya W, Glogauer M, Grinstein S. The phosphatidylserine receptor TIM4 utilizes integrins as coreceptors to effect phagocytosis. Mol Biol Cell 2014;25:1511–22.

51. Subramanian S, Parthasarathy R, Sen S, Boder ET, Discher DE. Species- and cell type-specific interactions between CD47 and human SIRPalpha. Blood 2006;107:2548–56.

52. Tsai RK, Discher DE. Inhibition of “self” engulfment through deactivation of myosin-II at the phagocytic synapse between human cells. J Cell Biol 2008;180:989–1003.

53. Kojima Y, Volkmer JP, McKenna K et al. CD47-blocking antibodies restore phagocytosis and prevent atherosclerosis. Nature 2016;536:86–90.

